# Tetraploid turnips (*Brassica rapa* ssp. *rapa*) are highly tolerant of tetravalent formation and aneuploidy

**DOI:** 10.1101/2025.06.04.657904

**Authors:** Zhenling Lv, Ingrid Schneider-Huether, Fei He, Annaliese S. Mason

**Affiliations:** Plant Breeding Department, INRES, University of Bonn, Kirschallee 1, 53115 Bonn, Germany; Department of Plant Breeding, Justus Liebig University, Heinrich-Buff-Ring 26-32, Giessen 35392, Germany

**Keywords:** meiosis, autopolyploid, *Brassica*, aneuploidy, tetravalents

## Abstract

Prior to 1980, experimental induction of polyploidy (chromosome doubling) led to the release of several tetraploid *Brassica rapa* ssp. *rapa* as fodder turnip cultivars. Most experimentally induced polyploids are meiotically unstable and show reduced fertility. However, we hypothesized that based on the requirement to produce large amounts of commercial seed to sell to farmers, these turnip lines should have managed to restore fertility and stabilize meiosis. We collected and tested all *B. rapa* listed as or referred to as tetraploid from the IPK Gatersleben, CGN Wageningen and Nordgen germplasm banks, and subsequently investigated chromosome karyotypes, meiotic chromosome behaviour and fertility in confirmed tetraploid turnip accessions, using a combination of resequencing and fluorescent in situ hybridisation. Contrary to our expectations, all accessions showed unstable meiosis: the average tetravalent frequency per meiosis per plant ranged from 4.8 to 6.4 per line, and these tetravalent associations also persisted from diakinesis to metaphase I. Using chromosome-specific fluorescence in situ hybridisation probes, we found that most chromosomes showed similar frequencies of tetravalent formation except for chromosomes A03 and A06, which predominantly formed tetravalents (>90%). Of the 21 individuals sequenced (one per accession), approximately half (9/21) were aneuploid (loss or gain of a whole chromosome), and two displayed additional chromosomal rearrangements. We nevertheless observed no significant phenotypic abnormalities or reductions in fertility (although all accessions were self-incompatible). Our findings indicate that stabilizing meiosis may not always be necessary to produce relatively fertile and homogeneous outcrossing polyploid populations.

## Introduction

Polyploidy refers to the condition in which an organism has more than two complete sets of chromosomes. Polyploid is widespread across many taxa, including insects, fish, algae, fungi and plants (Mable 2004). In plants, polyploidy plays a crucial role in evolution and speciation, often leading to increased genetic diversity, hybrid vigor, and improved adaptability to environmental conditions (Van de Peer et al. 2017; Heslop-Harrison et al. 2023). Autopolyploids result from genome duplication within a single species and lead to multiple sets of structurally similar homologous chromosomes (e.g. AAAA), in contrast to allopolyploids, which arise through hybridization between different species accompanied by the doubling of non-homologous (homoeologous) genomes (e.g. AABB) (Soltis et al. 2007; Barker et al. 2016). Despite this, novel autopolyploids must overcome a number of barriers to establish successfully. The most important barrier is quite probably regulation of meiosis, such that chromosome segregation occurs normally despite the presence of two very similar genomes (Cifuentes et al. 2010; Pelé et al. 2018).

The *Brassica* genus encompasses the widest variety of crop types of any plant genus, including cabbage, cauliflower, broccoli, swede (Rutabaga), kohlrabi, bok choy, pak choy, mustards, rapeseed, oilseed rape, canola, and more (Al-Khayri et al. 2021; Elena Cartea et al. 2021). As this group contains many vegetable types, the possibility to produce autopolyploids, which commonly have increased tissue and organ sizes (Doyle & Coate, 2019), offers high potential for yield improvement. The turnip (*Brassica rapa*) is an important vegetable crop, and valued for its versatility and nutritional benefits (Bradshaw et al. 2002; Abdel-Razzak 2021). Prior to 1980 there were many induced polyploidy experiments which led to the production and testing of different polyploid vegetable cultivars (Gowers 1977; Olsson and Ellerstrom 1981). In the 1950s-70s, mixed reports on the success of tetraploid turnip types (4*x Brassica rapa var. rapa*) were reported in France, Sweden and the Netherlands. Several of these experimental polyploid lines were subsequently released as cultivars, particularly as fodder turnips (Olsson and Ellerstrom, 1981), and autotetraploid turnip cultivars are still being grown today and are competitive with their diploid counterparts (Gowers, 1977; Abel & Becker, 2007; Meng et al., 2011).

In *Brassica*, induced or neopolyploidy is known to be associated with high levels of multivalent formation during meiosis (Cifuentes et al. 2010; Pelé et al. 2017). Formation of multivalents, which often leads to the production of gametes with aneuploid chromosome numbers, is thought to be a major cause of reduced fertility in autopolyploids (Darlington 1937; Birchler 2013). This irregular chromosomal segregation during meiosis results in unbalanced gametes, which can impair successful reproduction (Parisod et al. 2010; Bomblies et al. 2016; Bomblies 2023; Heslop-Harrison et al. 2023). Several studies on induced *B. rapa* (*B. campestris*) autotetraploids have showed varying levels of tetravalent formation during meiosis (Ramanujam and Deshmukh 1945; Swaminathan and Sulbha 1959). Over several generations of selection for fertility examined in one tetraploid genotype (1, 2, 17, 18 and 19 generations after induced polyploidization), bivalent formation increased and multivalent formation decreased, leading to improved seed fertility from 1.5 to 16.8 seeds per silique (Swaminathan and Sulbha 1959). However, the fundamental question of how exactly breeders managed to restore fertility in tetraploid lines in order to develop them as cultivars (e.g. as fodder turnips, which are propagated by seed) is still unknown. We hypothesized that, given the necessity of producing large quantities of commercial seed for farmers, these lines likely restored fertility and stabilized meiosis. This would make them an ideal model for studying the genetic mechanisms behind this restoration and exploring how autopolyploid crops could be successfully developed in the future.

In this study, we used reports of released tetraploid turnip cultivars from the literature (Olsson and Ellerstrom 1981, Gowers 1977) as well as accession record information to identify putatively tetraploid turnip accessions present in the IPK Gatersleben, CGN Wageningen and Nordgen germplasm banks as well as commercially available material. We subsequently identified and characterised 20 tetraploid turnip accessions from the germplasm banks and one commercially available line from seed company DSV (https://www.dsv-seeds.com/sorte/4572) using a combination of genome sequencing and cytogenetics analysis to investigate meiotic chromosome behaviour and chromosome number using fluorescent in situ hybridization (FISH).

## Materials and Methods

### Plant material

Detailed information on the plant material can be found in **Table S1 and Fig. S1**. Briefly, turnip accessions identified as putatively tetraploid through a combination of literature review and available passport information were obtained from three germplasm banks: IPK Gatersleben, CGN Wageningen, and Nordgen (M1 – 21), as well as additional turnip lines randomly selected as putative diploid controls, of which most were later also found to be tetraploid (M22 - M27; M28 was a confirmed diploid). An additional tetraploid line (M0) was sourced directly from DSV Seeds, a commercial seed company. Seeds from accessions M1 - M21 were germinated: three replicates from each accession were sown in quick-pots and seedlings transferred to 6 × 6 × 6.5 cm pots after 5 weeks for around 12 weeks of vernalization at 4 degrees C, followed by transplanting to 14.5 × 14.5 × 22.0 cm pots under glasshouse conditions at Justus Liebig University (JLU), Giessen, Germany. Seed setting data of accessions M0 and M22-M28 was generated later following the same procedure: germination, vernalization, and finally transplantation into 7.5 litre pots (approximately 27.5 cm diameter / 21.5 cm deep) in the greenhouse at the University of Bonn, Bonn, Germany.

### Cytogenetic analysis

We obtained meiotic chromosomes spreads which were subsequently used for Fluorescent in situ Hybridisation (FISH) following the protocols detailed in (Xiong and Chris Pires 2011; Cao et al. 2023), with minor modifications. Briefly, young floral buds were collected and fixed into Carnoy’s solution (3:1 ethanol: glacial acetic acid) for 12 hours at room temperature and then transferred to 70% ethanol at 4 °C for storage until use. The anthers at the right stage were collected and incubated in enzyme mixture (2% Onozuka R-10 cellulase + 1% Y23 pectolyase in 1 × citrate solution) for 90 minutes at 37 °C. Enzyme solution was rinsed away by cold 1 × TE, and TE was removed and anthers rinsed again in 100% ethanol several times (to ensure that all the water was eliminated). Anthers were then broken apart with a dull dissecting needle. 100% acetic acid was then added to increase the spread distance, following which 7 µl of cell suspension was dropped onto a glass slide. Meiotic chromosome spreads at diakinesis, metaphase I and anaphase I were used for FISH. Oligos for CentBr (*Brassica napus* centromeric repeat mixed probes CentBr1 and CentBr2) (Xiong and Chris Pires 2011), 5S (Waminal et al. 2018), and BAC KBrB072L17 (Xiong and Chris Pires 2011; Cao et al. 2023) were used for hybridisation; these probes contain repetitive elements or sequences which enable the identification of specific chromosomes. Detailed sequences and corresponding fluorophores ordered from Sigma-Aldrich can be found in Table S2. After hybridisation and washing of the slides, a drop of Vectashield Antifade Mounting Medium with DAPI (H-1200, Vector Laboratories) was added and then covered with a cover slip. Images were captured using the Zeiss Axio Observer 7 microscope equipped with Axiocam 305 camera. Contrast optimisation and image analysis was done using the Zeiss ZEN 3.4. software, Adobe Photoshop Elements software (v. 24.0, Adobe Systems Incorporated), and image sizes were adjusted in Microsoft PowerPoint 2019 (Microsoft Corporation).

### DNA extraction, sequencing and copy number variation analysis

DNA was extracted from young leaf samples at the 4 to 6 leaf stage of plant development using the Doyle and Doyle DNA extraction protocol (Doyle and Doyle 1987). Illumina paired-end sequencing was performed at Novogene Company Limited, United Kingdom on an Illumina HiSeq machine to produce paired end reads of 150 bp length. Clean reads were then mapped to the reference genome *B. rapa* ‘Chiifu-401-42’ version 4.1 (Zhang et al. 2023) using BWA Default parameter (Li and Durbin 2009). Removal of duplicates, sorting and indexing was carried out with Samtools (Danecek et al. 2021). SNP calling was performed using bcftools mpileup and filtered for a minimum quality of 30 and a minimum read depth of 10 using vcftools (Danecek et al. 2011). The phylogenetic tree was constructed based on a final dataset comprising a total of 3,454,715 snps using the Neighbor-Joining method by MEGA11 (Tamura et al. 2021). Specific chromosome copy number variants were identified based on normalised read coverage depth over 25 gene windows along the length of each *B. rapa* chromosome (Katche et al. 2023). For chromosomes with 4 copies, the normalized value is 1 (meaning that each individual chromosome corresponds to 0.25 after normalization). When an extra copy of a chromosome (5 copies) is present, the normalized value would be 1.25, while a normalized value of 0.75 means loss of one chromosome copy (3 copies).

### Fertility assessment

Pollen viability and number of seeds produced per ten pods were scored to describe fertility. Pollen viability for accessions M1-M21 at Justus Liebig University (JLU) was assessed for two freshly opened flowers per plant, and pollen grains stained with 1-2% acetocarmine solution (Leflon et al., 2006). At least 600 pollen grains per plant were counted and pollen viability was assessed using the Zeiss Axio Observer 7 microscope, equipped with Axiocam 305 camera, and the images were analysed using a Zeiss Zen v3.4 microscope (Carl Zeiss, Oberkochen, Germany) assuming darkly stained (red) pollen grains were viable, and weakly stained or shrivelled pollen grains were non-viable. For the seed number, we failed to notice the self-incompatibility of the turnip plants during the first year of the experiment (2019-2020), and subsequently did not obtain any seeds or only obtained a very small number per plant. To address this issue, we conducted a follow-up experiment in which 4–5 plants for accessions M0 and M22-M28 at the University of Bonn were grouped together for cultivation (2022-2023). Upon harvest, we successfully obtained pods with seed. The number of seeds produced per ten pods was counted for each plant after harvesting.

### Statistical analysis

Statistical analyses were carried out using RStudio version 4.0.2 (2020-06-22). Figures were generated in RStudio and edited with PowerPoint 2019 (Microsoft Corporation).

## Results

### Tetravalent and bivalent configurations identified by FISH in tetraploid turnip accessions

We examined chromosome configuration at diakinesis, as chromosomes are more dispersed during this stage, making observation and analysis more convenient. To aid in identifying and analyzing the chromosomes, which are small and undifferentiated in *Brassica*, we utilized a set of chromosome probes: centromere probes (CentBr1+ CentBr2) designated as Oligo-CentBr, Oligo-5S and Oligo-BAC KBrB072L17 (**Fig. 1**). 5S signals were observed on chromosomes A01, A03 and A10 with the strongest 5S signal located in the pericentromeric region of chromosome A01. In addition, A01 showed oligo-BAC KBrB072L17 signals at the top of its short arm, while A03 displayed 5S signals without any corresponding oligo-BAC KBrB072L17 signals. A10 also exhibited 5S signals and is the smallest A-genome chromosome. oligo-BAC KBrB072L17 signals appeared on A05, A04, A02, and A06. Of these, A05 displayed the strongest oligo-BAC KBrB072L17 signals. Chromosome A04 is smaller than both A02 and A06, with A06 showing stronger oligo-BAC KBrB072L17 signals than A02. For the remaining chromosomes, A07, A08, and A09, A08 is the smallest of the three, A09 exhibited faint oligo-BAC KBrB072L17 signals under overexposure, and no signals were detected on A07 (**Fig. 1**). This method is both highly effective and efficient. While DAPI staining can be used to identify tetravalents, it can be challenging in *Brassica* species, where small chromosomes make it difficult to distinguish between the tetravalents of small chromosomes and the bivalents of larger ones. However, by using this probe combination, we can clearly differentiate between a bivalent formed by two chromosomes and a tetravalent formed by four chromosomes, allowing for accurate and efficient identification of tetravalents.

**Fig. 1:**
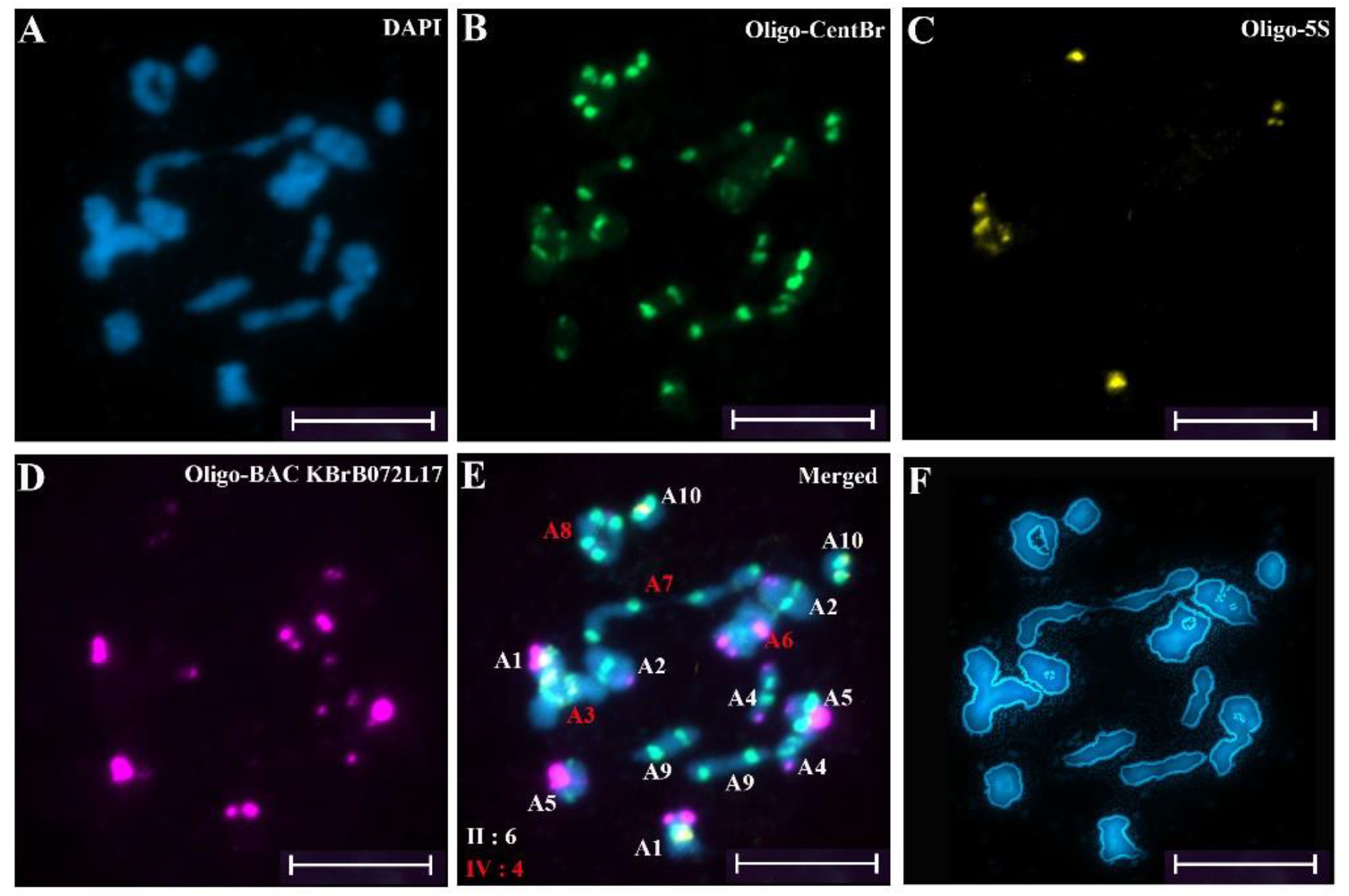
Characterization and quantitative analysis of chromosome configurations. A-E Diakinesis in an autotetraploid turnip plant using A: DAPI; B: Oligo-CentBr; C: Oligo-5S; D: Oligo-BAC KBrB072L17 to identify individual chromosomes; E: Merged fluorescence image with specific chromosomes labelled, showing chromosome configurations: A03, A06, A07 and A08 as tetravalents and A01, A02, A04, A05, A09 and A10 as bivalents. F: clear display of chromosome configuration outlines. This image example originated from the M15 accession. Scale bar represents 10 µm.

### Tetravalents are common in tetraploid turnip accessions

Using this method, we were able to easily identify and distinguish tetravalents. During the analysis of tetravalent configurations (representative, well-defined, and distinguishable tetraploid configurations isolated from a variety of distinct cell types are presented), we found that chromosome configurations at diakinesis were primarily observed as ring and chain structures (**Fig. S2**). We observed that, across all materials analyzed, an average of 4.6 to 6.4 tetravalents were formed per accession (**Fig. 2A**, detailed information including bivalent number, tetravalent number, and cell numbers counted is summarized in **Table S3**). We exclusively selected euploid materials for cytological analysis to avoid potential interference from aneuploidy or other confounding factors. Among the 9 euploid accessions we analyzed, the lowest average number of tetravalents was observed in M17 (4.6), while the highest was found in M19 (6.4). A one-way ANOVA test revealed significant differences between accessions. Nevertheless, all accessions, each possessing the basic chromosome number of 10, consistently exhibited 4 to 6 tetravalents on average per PMC.

**Fig. 2:**
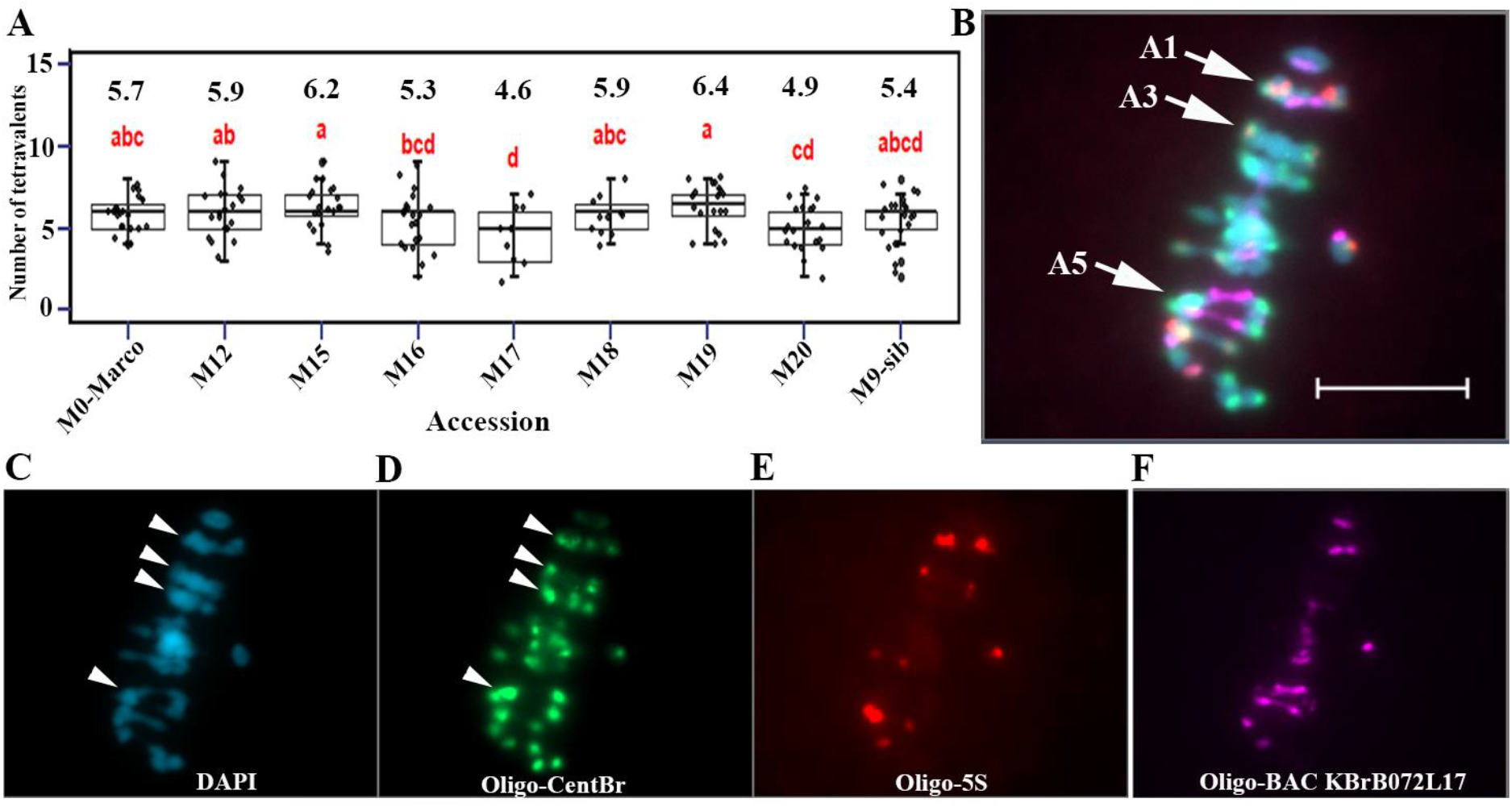
Diakinesis tetravalent configurations are common in tetraploid turnip accessions and persist to metaphase I. A: significant differences were observed between lines (one-way ANOVA, p = 0.0088); different letters indicate significant differences (LSD post-hoc test, p < 0.05); B: tetravalent configurations for A01, A03 and A05 (white arrows) observed in metaphase I; C: DAPI; D: Oligo-CentBr; E: Oligo-5S; F: Oligo-BAC KBrB072L17 fluorescence image to recognize chromosomes, A1, A3 and A5 can be easily recognized and the other tetravalent would be A7 or A9. This image example originated from the M0-Marco accession. Scale bar represents 10 µm.

Tetravalents were also observed during metaphase I, suggesting tetravalents persist from diakinesis until metaphase I without being resolved (**Fig. 2B, and Fig. 2C-2F** representing corresponding fluorescence channels). We also conducted a quantitative cytological analysis (**Table S4**); however, the number of cells which were observed was very limited. Furthermore, overlapping cells during metaphase I of meiosis posed challenges for accurate observation. Despite these limitations, we still detected an average of 2.3 tetravalents at metaphase I. Due to low cell numbers we were unable to conclude if this was significantly different from diakinesis tetravalent frequencies (whether some tetravalents might be resolved), but we could establish that tetravalents were definitely still persisting from diakinesis to metaphase I.

In addition, we not only identified tetravalents but also observed univalents that fail to pair with their homologous counterpart, with no visible physical linkage (**Fig. 3A and Table S3**). Interestingly, most chromosomes had similar likelihoods of forming bivalents vs. tetravalents (33.5 - 66.5%), with the exception of chromosomes A03 and A06 which showed a >86% tendency to form tetravalents (Bimodal Distribution test with p-value < 2.2e−16). Additionally, the formation of univalents displayed a distinct pattern, predominantly occurring for chromosomes A10, A08, A09, A07 and A04 (one-way ANOVA, p < 2.2e−16) (**Fig. 3B and Table S3**). Among the analyzed materials, A10 exhibited the highest frequency of univalent formation, with approximately 5.5% of examined cells showing univalent formation of A10, followed by A8 at 4.8% and A4 at 3.6%. The presence of univalents in these tetraploids further increases the risk of chromosome mis-segregation during the later stages of meiosis.

**Fig. 3:**
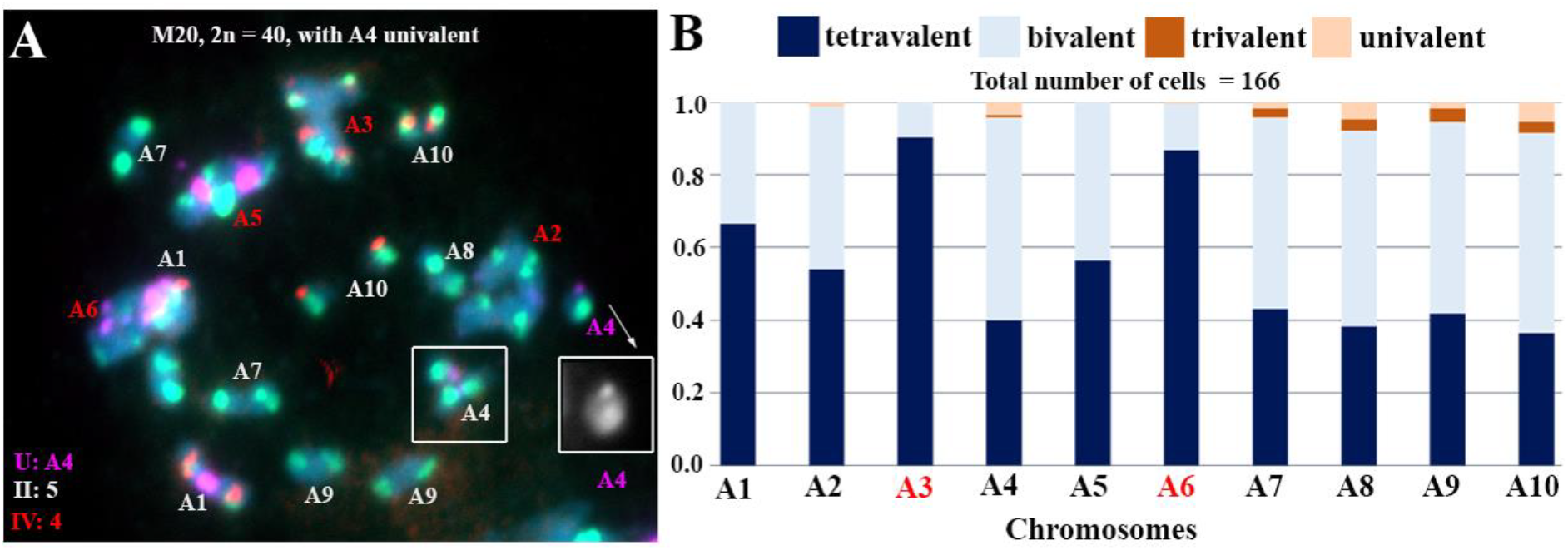
Distribution proportions of chromosomal configurations across different chromosomes. A: typical univalent structure for chromosome A04, indicated by white box; B: the frequency of different chromosome configurations: red font indicates chromosomes A03 and A06 which have a significant tendency to form tetravalents. This image example originated from the M20 accession. Scale bar represents 10 µm.

### Aneuploidy was common in autotetraploid turnip lines

Given that these materials are commercial varieties capable of producing a large number of seeds and are theoretically expected to be genetically stable, we randomly selected one individual from each of the 21 tetraploid turnip lines (**Fig. 4A**) for resequencing and genomic analysis. Considering that these materials are tetraploid, we performed DNA resequencing at a depth of 30×, ensuring sufficient data for phylogenetic analysis (SNPs) and chromosome copy number assessment (see methods section). Based on the phylogenetic tree, there was little observable genetic structure across the accessions, despite variation in country of origin (Germany, Netherlands, Poland, Sweden and Denmark) and collection year (1960 - 2021) (**Fig. 4B and Table S5**). We also discovered that 9/21 plants were aneuploid (loss or gain of whole chromosomes) for different chromosomes: M4, 2n = 39, loss of A01; M6, 2n = 39, loss of A07; M7, 2n = 39, loss of A08; M3, 2n = 40, loss of A01 and gain of A07; M8, 2n = 41, gain of A04; M5, 2n = 41, gain of A09; M9, 2n = 41, gain of A10; M11, 2n = 42, gain of A2 and A10 (**Fig. 3C**). Our observations of three or five copies of individual chromosomes suggests that these aneuploidy events have not been fixed and will continue to segregate in subsequent generations.

**Fig. 4:**
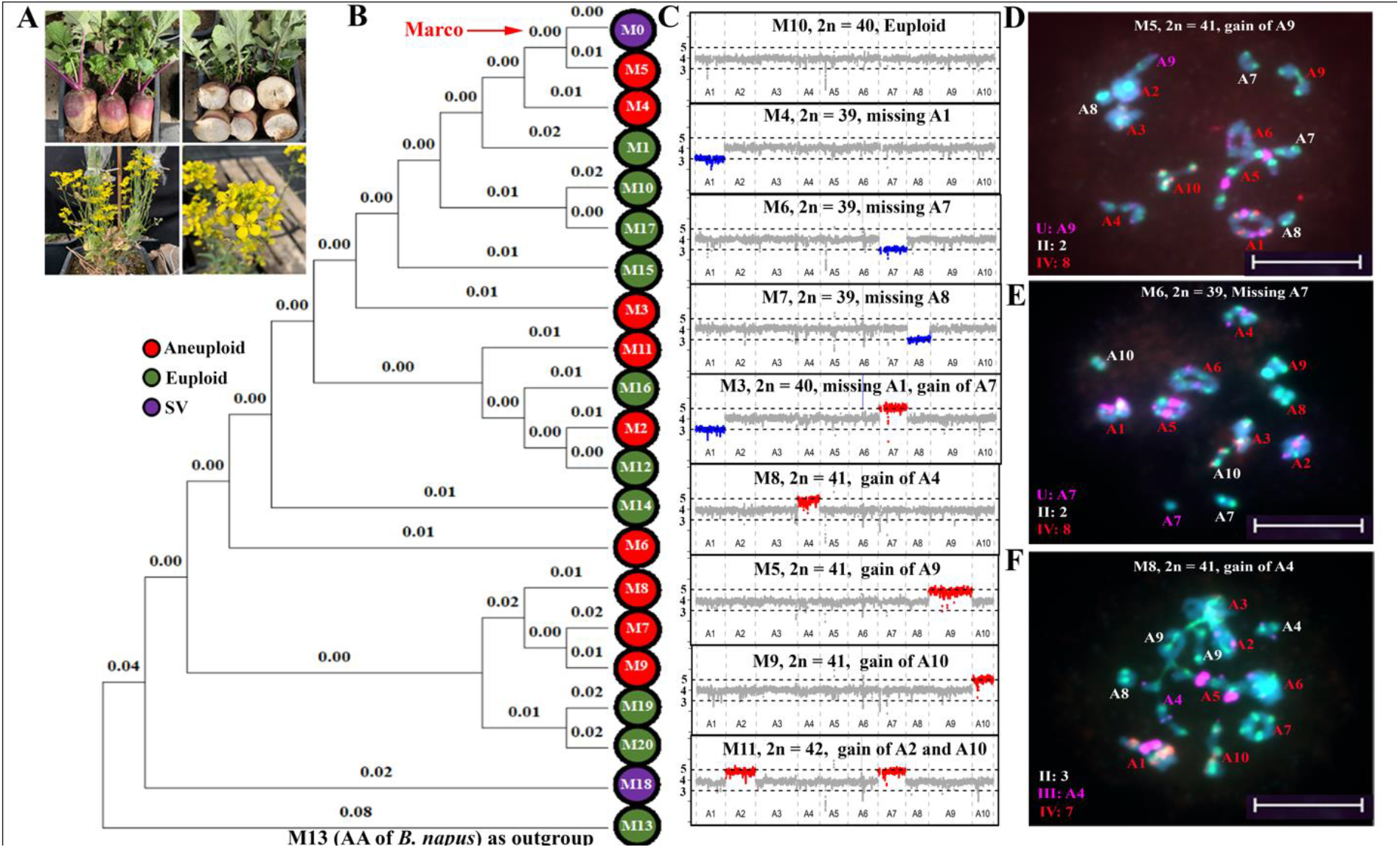
Aneuploidy was common in autotetraploid turnip lines. A: phenotype of root tuber turnip commercial line M0 “Marco”; B: the dendrogram generated by MEGA11 software using 3 454 715 SNPs obtained from sequencing data indicating genetic differences between samples; C: normalised read coverage depth derived from 30 × sequence coverage per sample over 25 gene windows across the ten *B. rapa* chromosomes. The dashed lines indicate an extra copy of a chromosome (5 copies), regions highlighted in red, and loss of one chromosome copy (3 copies), regions highlighted in blue; D-F: cytological images confirmed the accuracy of the sequencing data: (D) M5, 2n = 41, gain of A09; (E) M6, 2n = 39, missing A07; and (F) M8, 2n = 41, gain of A04. Scale bar represents 10 µm.

We subsequently confirmed the reliability of our sequence data analysis through cytological observation. Using the same plant whose leaves were used for resequencing, meiosis was observed from flower buds, and in all cases confirmed the sequence-based predictions (**Table S1**). We found that in a single meiotic cell of M5, 2n = 41, five chromosome A09 copies were present, and formed one tetravalent and one univalent in the given cell (**Fig. 4D**); that in M6, 2n = 39, three copies of A07 were present and formed one bivalent and one univalent (**Fig. 4E**); and in M8, 2n = 41, five copies of A04 were present and formed one bivalent and one trivalent (**Fig. 4F**).

To determine whether the siblings of these aneuploids were euploid or not, we used the same cytological methods to examine different individual plants from several accessions. Additional plants of M7 and M9 were both euploid, in contrast to their aneuploid siblings, while a sibling of M3 had 2n = 38, missing copies of A07 and A05 - different aneuploidy to the sequenced plant which had a missing copy of A01 and an extra copy of A07 (**Fig. S3**).

### Chromosomal structural rearrangements were also observed

Based on our resequencing results, we identified chromosome structural variation (involving duplication or deletion of one or more chromosome segments) in two samples, M18 and M0 (Marco), and we used the same individual plants which were sequenced to examine the chromosome karyotypes cytologically (**Fig. 5**). In M18, three chromosomes - A02, A06, and A08 - exhibited structural changes (**Fig. 5A**). The top of chromosome A02 from 0 - 4 Mb was present in only two copies, while the segment from 4 - 25 Mb was present in 3 copies. Cytological observation revealed a translocation event between chromosomes A04 and A02 (**Fig. 5A**). Furthermore, we observed the same A04 – A02 translocation in a sibling of M18 (**Fig. S4**). Chromosome A06 was also confirmed to be missing a segment, although this did not appear to impact chromosome pairing and quadrivalent formation during metaphase I (**Fig. 5C**). The interstitial duplication present for part of chromosome A8 could not be definitively observed cytogenetically (**Fig. 5C**).

**Fig. 5:**
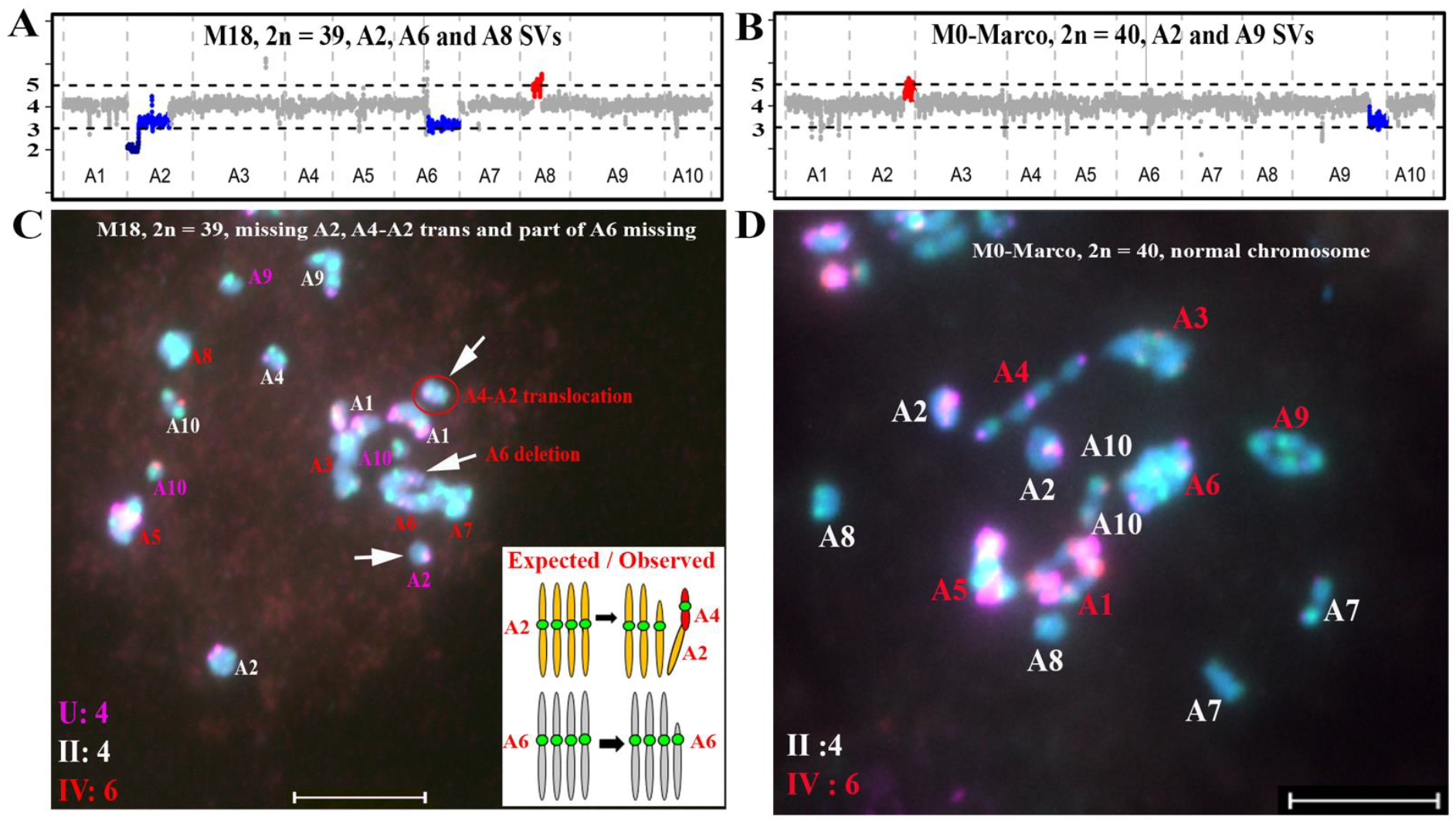
Chromosomal translocation detection by FISH. A and B: copy number variation via normalized read coverage depth over 25 gene windows across the ten *B. rapa* chromosomes in M18 and M0-Marco. The dashed lines indicate an extra copy of a chromosome (5 copies, red color) and loss of one chromosome copy (3 copies, blue color). C: diakinesis cell showing (indicated by arrows) a truncated A02, an A04-A02 translocation, and a missing chromosome segment in the A06 tetravalent configuration in M18; and D: normal diakinesis cell in M0-Marco, no obvious chromosomal abnormalities are visible. Scale bar represents 10 µm.

In M0 (Marco), we detected a terminal deletion on chromosome A09 (three copies) and a terminal duplication on chromosome A02 (five copies) (**Fig. 5B**). The odd number of chromosome copies involved indicates that this chromosomal variation is not fixed, and is most likely novel and still segregating. Due to the small size of this region and the coverage of the FISH markers, we were unable to determine whether this deletion is due to a translocation or not (**Fig. 5B**).

### Fertility is quite normal in commercial turnip lines

Pollen fertility was generally high across all plants regardless of euploid or aneuploid status, with averages exceeding 85%, although M18 (one of the two lines with structural variation) showed slightly lower fertility than the other lines (**Table S6 and Fig. 6**). Pollen viability was not significantly different between aneuploid and euploid lines (sequenced lines, Fig. 6). No plants from accessions M1 – M21 set self-pollinated seed when bagged (under glasshouse conditions in 2020), indicative of self-incompatibility.

**Fig. 6:**
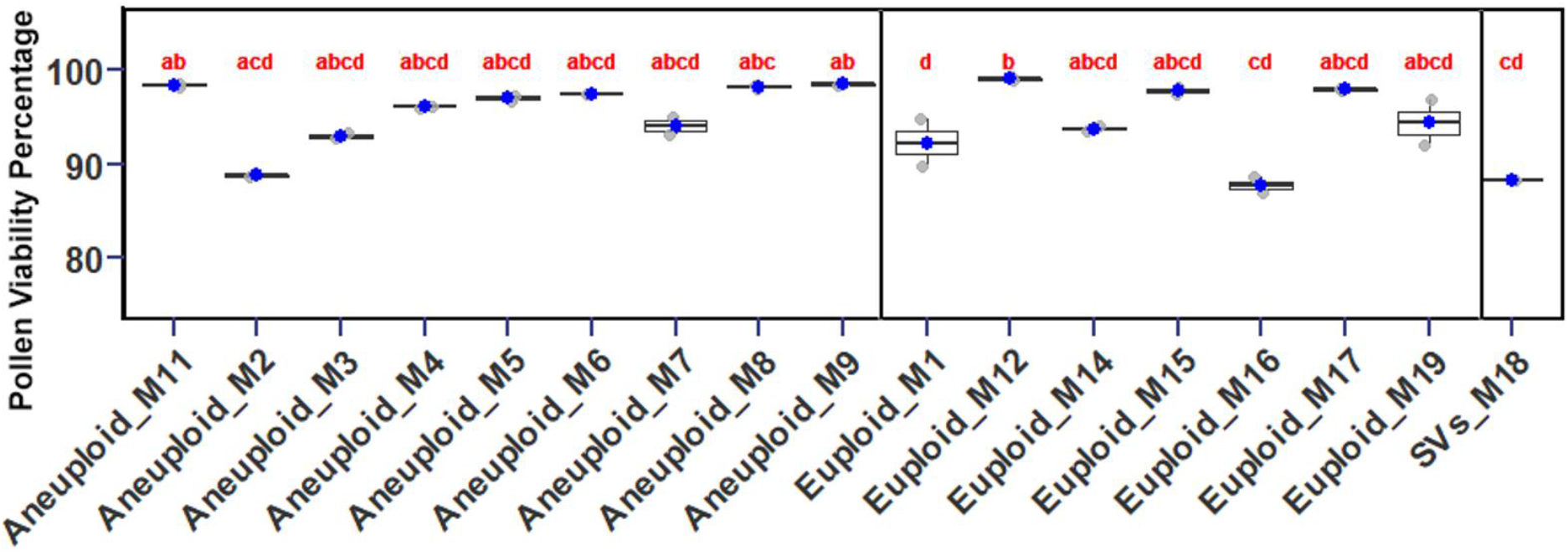
High pollen viability was observed in both euploids (2n = 40) and aneuploids of tetraploid turnip. Significant differences were observed between lines (Kruskal-Wallis test, p = 0.0083); different letters indicate significant differences (Dunn post-hoc test, p < 0.01).

To confirm that the tetraploid turnips were in fact able to set seed under open pollination conditions, we investigated fertility in a further eight lines which were available and already through the 12 week vernalization period which we had also found to be tetraploid (M0, M22-M27, detailed information on the plant material can be found in **Table S1**) by enclosing 4-5 plants with plastic sheeting to facilitate cross-pollination and counted the number of seeds per 10 pods for each plant (**Table S7**), using a diploid turnip line (M28) as a control (**Fig. S5A**). Despite some differences between genotypes for seed set, we observed no significant difference in seed set between the tetraploid lines and the diploid fodder turnip control line (**Fig. S5B and Table S7**).

## Discussion

### Tetravalents are prevalent in tetraploid turnip meiosis

We observed high frequencies of tetravalents in our tetraploid turnip lines. Cytological studies have consistently found that tetravalent formation frequently occurs during autopolyploid meiosis in both naturally established species and synthetic lines across various plant taxa (reviewed by (Lv et al. 2024)), including kiwifruit (Wu et al. 2014), *Brassica* (Howard 1939; Swaminathan and Sulbha 1959; Zdráhalová 1968; Jenczewski et al. 2002), cereals and grasses (Morrison and Rajhathy 1960 a, b), and potato (Choudhary et al. 2020). In synthetic autopolyploids, frequent multivalents are expected, often leading to fertility issues, while in naturally established autopolyploids, evolutionary selection is thought to lead to a more diploidized meiosis with predominantly bivalent formation (Ramsey and Schemske 2002; Soltis et al. 2007; Parisod et al. 2010), as is observed in established autopolyploid *Arabidopsis arenosa* (Bomblies et al. 2016; Morgan et al. 2021). Hence, we also hypothesized that the established autopolyploid turnip lines would show diploidized meiosis, as a result of agricultural selection for seed fertility. However, the average tetravalent frequency per meiosis per plant ranged from 4.8 to 6.4 per line, and these tetravalent associations also persisted from diakinesis to metaphase I and were not resolved, disproving our hypothesis. During the examination of tetravalent frequencies, we observed that chromosome configurations at diakinesis were predominantly ring and chain structures, which often resulted in equal distribution and segregation of tetravalents to opposite poles during anaphase I. Parallel and convergent centromere orientations can lead to a balanced, half-and-half distribution of quadrivalents to the poles (Rieger et al. 1976; Sybenga 1995). However, we did also see frequent aneuploidy, suggesting that not all tetravalents resulted in balanced chromosome segregation. Regardless, from our results it is evident that tetravalent formation was not as detrimental as we may have expected in these lines.

### Tetravalent frequencies differed by A-genome chromosome

On average, all individual A-genome chromosome sets formed tetravalents at 33 – 67% frequency, except for chromosomes A03 and A06 (>86% frequency). Crossover frequency per chromosome in *Brassica* is known to be correlated with chromosome length (Mason *et al*., 2016; Higgins *et al*., 2018). As A03 is the largest A-genome chromosome (Xiong *et al*., 2011), this may partially explain why it has more tetravalent formation (as tetravalents are more likely to form with additional crossovers). However, A09 is the second largest chromosome and did not show increased tetravalent frequency, whereas A06 is intermediate in size and frequently showed tetravalents. Further investigation of crossover distribution frequencies and structural features of these chromosomes might shed light on these results. Notably, A6 was previously reported by Xiong et al. (2011) to exhibit the lowest rate of chromosomal variation in the A genome, which may suggest the presence of unique sequences or structural features. These findings highlight the need for further investigation into specific chromosome features in future studies.

### Aneuploidy was common in tetraploid turnip plants but not fixed

Loss and gain of individual chromosomes (aneuploidy) was common in all the autotetraploid turnip lines investigated, but no individuals showed loss or gain of more than one copy of a homologous chromosome. Of the 21 individuals sequenced, approximately half (9/21) were aneuploid (loss or gain of a whole chromosome), and further cytological observations of siblings within the same line as sequenced euploid individuals revealed ongoing karyotype instability. These findings suggest that none of these commercial turnip lines are genomically stable. Surprisingly though, it also seems that karyotype variation is not accumulating over generations in these lines, as has been observed in other meiotically unstable *Brassica* systems (Mwathi et al. 2019). Possibly, as commercial turnips reproduce through traditional open pollination, this process promotes not only greater genetic diversity (Bradshaw et al. 2002) but reduces the likelihood of stabilizing or fixing chromosomal aneuploidy. In self-pollinating species, chromosomal imbalances are more likely to become fixed over generations, as progeny are expected to segregate for single-copy chromosomes in a 1 : 2 : 1 ratio of 2 copies : 1 copy : 0 copies of the chromosome, where 2 or 0 copies are then stably inherited in the following generation (i.e. an aneuploid karyotype may become “fixed” as two rather than four copies of a single chromosome). However, in outcrossing populations, assuming that chromosome aneuploidy is evenly distributed over the genome, as appears to be the case here, the likelihood that two individuals have the same missing or additional chromosomes is reduced, lowering the risk of inheriting or perpetuating aneuploidy and resulting in healthier offspring with balanced genomes. Due to the limitations of our current data, we were unable to determine whether any particular chromosome was more likely to be lost or gained: this would need to be investigated in much larger segregating populations in future.. Whether or not selection (e.g. at the embryo or seed development level against more serious chromosomal imbalances) plays a role in this effect would also be an interesting topic for further investigation.

### Aneuploidy had no obvious impact on fertility or plant appearance

Surprisingly, we saw no major impact of aneuploidy on fertility or on qualitative appearance of the plants: although we did not originally expect aneuploidy or phenotype for specific differences between plants, fertility data and appearance were not clearly different between euploid and aneuploid tetraploid turnip plants. The limited impact of aneuploidy on phenotype in our study could be explained by gene expression regulation, as is the case with wheat and autotetraploid potato (Stupar et al. 2007; Zhang et al. 2017; Zeng et al. 2020): perhaps having three or four copies of a gene does not significantly alter overall expression levels, mitigating the potential for harmful effects. In yeast, over 70% of proteins encoded on aneuploid chromosomes were found to undergo dosage compensation via protein turnover, aligning their average protein levels more closely with the euploid state (Muenzner et al. 2024). Together, transcriptomic and proteomic studies have provided varying levels of insight into the buffering capacity of polyploid genomes, and would be interesting to explore also in autotetraploid turnips.

### Chromosome rearrangements were found in a current commercial line

Another surprising finding was the observation of structural chromosome variation in M18 and M0-Marco. Our results suggest that chromosome segment duplications and deletions may be more frequent than previously thought in autopolyploids (in our case 2/21 plants). While karyotype rearrangements occur over time in most species, they likely happen at very low frequencies (Weiss and Maluszynska 2001; Parra-Nunez et al. 2019; Mengist et al. 2023). However, *Brassica rapa* is also a mesopolyploid, with a triplicated genome structure as a result of two rounds of ancestral polyploidy events (Parkin et al. 1995; Parkin et al. 2003; Cheng et al. 2014), which could also explain this higher-than-expected frequency of autosyndetic pairing. Autosyndesis has also been observed in haploid *B. rapa* (up to 3 bivalents per cell, Armstrong & Keller, 1981) and detected in *Brassica* interspecific hybrids with a haploid A genome (Mason et al. 2010; Mason et al. 2014). Such pairing disruptions may further contribute to chromosome variation and karyotype changes, including the A04-A02 translocation, A09 terminal deletion, and A02 terminal duplication we observed. Possibly, looser selection against such chromosomal rearrangements could exist because of the robust genomic buffering capacity provided by multiple sets of chromosomes at the autotetraploid level: polyploidy might reduce the lethality of such events and mitigate the consequences of these rearrangements (Parisod et al. 2010; Hollister 2015; Van de Peer et al. 2017).

### Fertility was not correlated with meiotic abnormalities

Although tetravalent formation and non-homologous chromosome interactions are generally thought to be highly detrimental to fertility, we found little correlation between meiotic behavior, chromosome complement, pollen fertility and seed setting in the tetraploid turnips. This might be due to the fact that fertility is not under strong selection in these *B. rapa* turnip plants (Bradshaw et al. 2002; Elena Cartea et al. 2021), as expected. Collecting fertility data in our study was also challenging because the plants are outcrossers, and the conditions under which the data were collected did not perfectly mimic field conditions. This makes it difficult to draw strong conclusions about the true levels of fertility under natural or field conditions. As an outcrossing root vegetable cultivar primarily used for animal feed, selection for traits such as yield, size, or disease resistance might be prioritized over seed fertility by breeders.

The frequent formation of tetravalents during meiosis also suggests that diploidized meiosis may not be under strong selective pressure in autotetraploid commercial turnips. This could be due to the relatively weak correlation between diploidized meiosis and fertility in these plants. In polyploids, where multiple chromosome sets are present, the occurrence of tetravalents—where four homologous chromosomes pair together during meiosis—does not necessarily result in reduced fertility. Unlike in diploid species, where precise pairing of homologous chromosomes is essential for successful gamete formation, polyploids often tolerate more complex meiotic configurations without significant reproductive consequences. Recently, non-meiotic factors such as pollen tube growth have also been shown to affect polyploid fertility in neopolyploid *A. arenosa* and *A. thaliana* polyploids (Westermann et al. 2024). Obviously, polyploid fertility is a complex phenomenon influenced by both genetic factors related to meiotic chromosome segregation and other contributing factors. The existence of other contributing factors to polyploid fertility (other than meiotic behaviour) was already proposed 75 years ago by G.L. Stebbins (Stebbins 1947), but what meiosis-independent additional factor or factors might influence fertility in tetraploid turnips continues to await further exploration and discovery.

## Author Contribution Statement

Z.L. performed experiments. ISM grew plants and collected samples. FH optimised the copy number variation analysis pipeline for the sequencing data. Z.L. and A.S.M. analysed, interpreted data, and drafted the manuscript. A.S.M. conceptualized and designed the project, acquired funding and supervised Z.L.. Z.L. and A.S.M. critically revised the manuscript. All authors read and approved the final manuscript for submission.

## Acknowledgements

We sincerely thank Prof. Zhiyong Xiong from Inner Mongolia University, China, for kindly providing us with the BAC-KBrBo72L17 oligo sequences. This project was funded by the German Research Council (DFG grant MA6473/12-1, awarded to AM) and by the European Union (ERC, SAMEY, 101087575). Views and opinions expressed are however those of the author(s) only and do not necessarily reflect those of the European Union or the European Research Council. Neither the European Union nor the granting authority can be held responsible for them. The Mason lab is partially funded by the Deutsche Forschungsgemeinschaft (DFG, German Research Foundation) under Germany’s Excellence Strategy – EXC 2070 – 390732324.

## Data Availability

The data that supports the findings of this study are available in the supplementary material of this article. The raw Illumina sequencing data of the 21 accessions are available at the NCBI Sequence Read Archive (SRA), under BioProject PRJNA1212698.

## Supplemental Tables

**Table S1**. Detailed information of 29 fodder turnip accessions.

**Table S2**. Oligo sequences of CentBr, 5S and BAC KBrB072L17 and corresponding fluorophores ordered from Sigma-Aldrich and used for hybridisation.

**Table S3**. Detailed scoring for chromosome configurations in euploid fodder turnip accessions.

Table S4. Detailed scoring for tetravalent configurations in metaphase I.

**Table S5**. Detailed statistics of resequencing data and read mapping, and unique read mapping ratio for all samples.

**Table S6**. Pollen viability assessment in two freshly opened flowers per tetraploid turnip plant.

**Table S7**. Number of seeds produced per ten pods counted for each tetraploid turnip plant after harvesting.

